# Long-term moisture barrier performance of liquid crystal polymer for implantable medical electronics

**DOI:** 10.64898/2026.02.24.707821

**Authors:** Brianna Thielen, Christopher Pulicken, Eyal Aklivanh, Philip Sabes, Milan Cvitkovic

**Affiliations:** Integral Neurotechnologies, South San Francisco, CA, USA

## Abstract

Liquid crystal polymer (LCP) is commonly used in the electronics industry due to its favorable dielectric, thermal, and insulative properties. It has recently gained popularity in the medical field for these same reasons, as well as its biocompatibility, moisture barrier properties, and ability to be microfabricated into thin film flexible circuits or flex PCBs. While polymers such as polyimide and Parylene C remain more common for electronics encapsulation and flexible circuit fabrication due to their relatively lower barriers to adoption and history of use, LCP’s superior moisture barrier performance and low risk of delamination make it a promising material for chronic use in medical devices. In this work, the moisture barrier properties of LCP are evaluated using *in vitro* accelerated aging over 59-61 weeks at 65-68 °C, corresponding to an equivalent implanted lifetime of 8.1 and 9.4 years at 37 °C for each of two sample groups: LCP as an electronics encapsulant and as a flexible circuit substrate. In the encapsulation group, relative humidity inside an encapsulation pocket was monitored over time with no noticeable change in humidity throughout the measurement period. In the flexible circuit group, impedance of laminated interdigitated electrodes was monitored over time, with an average decrease to 44% of the initial impedance value across all successful samples due to the moisture absorption of the LCP, which has remained stable for the latter half of testing. In both groups, no delamination was observed. These findings demonstrate that LCP is a viable moisture barrier for electronics in implanted medical devices for an estimated equivalent lifetime of at least 8.1 years.

## 2. Background

Liquid crystal polymer (LCP) is a promising material for the fabrication of thin film electrodes, the encapsulation of medical implants, and the fabrication or protection of electronics in harsh environments. While its most common use is in injection molded housing of electronic components, it also has a significant use in the medical field, owing to its USP class VI biocompatibility and advantageous properties over many other medical polymers [1]. For medical devices requiring contact with wet environments, LCP has a very low water vapor transmission rate (WVTR) and water absorption [2,3], allowing it to serve as a protective layer and maintain geometric stability over time. For devices which require high temperature fabrication steps, LCP has a high melting temperature, low coefficient of thermal expansion [4,5], and its liquid crystal property allows it to withstand many heat cycles without significantly altering mechanical properties.

While LCP is more commonly used as a structural material in existing medical devices [1], it also has very promising potential as a dielectric material in flexible electronics for medical applications. Flex PCBs can be fabricated using a high melting temperature LCP in place of polyimide and a low melting temperature LCP in place of the coverlay, bonding all layers together with thermal lamination [6]. LCP can also be used to produce microfabricated thin film polymer devices [2] in place of traditional thin film polymers such as polyimide or Parylene C, although some constraints arise in the availability of LCP film thicknesses (25, 50, 75, and 100 μm standard thicknesses) and the need for processing equipment not found in most cleanrooms (high temperature lamination, laser etching).

LCP also shows significant promise in the protection of electronic components from moisture. This is particularly useful for components populated on LCP flexible circuits, where an LCP “:cap”: can be formed and sealed over the component in place, protecting it from the surrounding environment [2,7], or in the creation of near-hermetic enclosures with electrical feedthroughs similar to hermetic titanium cans often used in medical implants [8].

When compared to similar devices built from other commonly used polymers (polyimide, Parylene C), LCP performs significantly better in applications requiring long term exposure to warm, wet environments, such as is required in a medical implant. There are two primary failure modes for polymer-encapsulated electronics in wet environments: moisture ingress through the polymer, and delamination of polymer-polymer interfaces. LCP has a WVTR of approximately 0.2 g/m^2^/day [3], as compared to 2-4 and 8 g/m^2^/day for polyimide [9] and parylene C^1^ [10], respectively, minimizing fluid transport through the polymer. LCP also has better inter-layer adhesion strength due to high chain mobility near its melting temperature and ability to crosslink between layers, minimizing the risk of delamination at polymer-polymer interfaces.

While these material properties are known and some studies have demonstrated LCP-based electrode longevity on a time scale of 2-3.5 years [3,11], there is limited data on the longevity of the material itself under simulated implanted conditions. This study investigates the chronic insulation performance of LCP as an encapsulant and a flexible circuit dielectric material in a simulated phosphate buffered saline (PBS) environment. The experiment was run at elevated temperature, equating to 8+ years equivalent implanted time to date, with the experiment ongoing. All data (as published here and ongoing) are publicly available on GitHub (see supplementary data).

## 3. Methods

LCP is primarily used as a moisture barrier material in one of two ways: in the form of an encapsulating pocket around electronic or other moisture sensitive components, or in the form of a laminated assembly, such as a flex PCB or microfabricated thin film device. In both cases, the two major failure modes are fluid transfer through the LCP (often appearing gradually over time) and delamination of LCP at sealed edges between two pieces of LCP (often appearing rapidly). Two sample groups were fabricated to test both of these barrier capabilities and failure modes.

### 3.1 Encapsulation Samples

To measure encapsulation robustness, LCP pockets were fabricated to encapsulate humidity sensors while submerged in 1x phosphate buffered saline (PBS). The relative humidity (RH) of the air inside the pocket was monitored.

Two humidity sensors were selected for encapsulation: a capacitive RH and temperature sensor system in package (CC2D25-SIP, Telaire, St. Marys, PA, USA) and a resistive RH sensor (HS30P, Telaire, St. Marys, PA, USA), referred to herein as the capacitive and resistive sensors, respectively. The capacitive sensor was selected for its smaller size and higher accuracy, while the resistive sensor was also included for its similarity in function to the second sample group (interdigitated electrodes). Pins of each sensor were soldered to 12”: hookup wire and insulated from each other using heat shrink. Header pins were soldered to the free end of the hookup wire.

For each sensor, a deep LCP pocket was fabricated using the process described in the following paragraph and illustrated in Figure 1 to seal the sensor on all but one side, where the sensor leads exited. Two sensors of each type were encapsulated in each of two LCP thicknesses (25 and 100 μm), producing 8 samples total. Pocket dimensions were 6 × 10 × 80 mm and 12 × 20 × 80 mm for the capacitive sensors and resistive sensors, respectively, with one of the small ends (6 × 10 mm and 12 × 20 mm) open for the sensor leads to exit.

**Figure 1.**
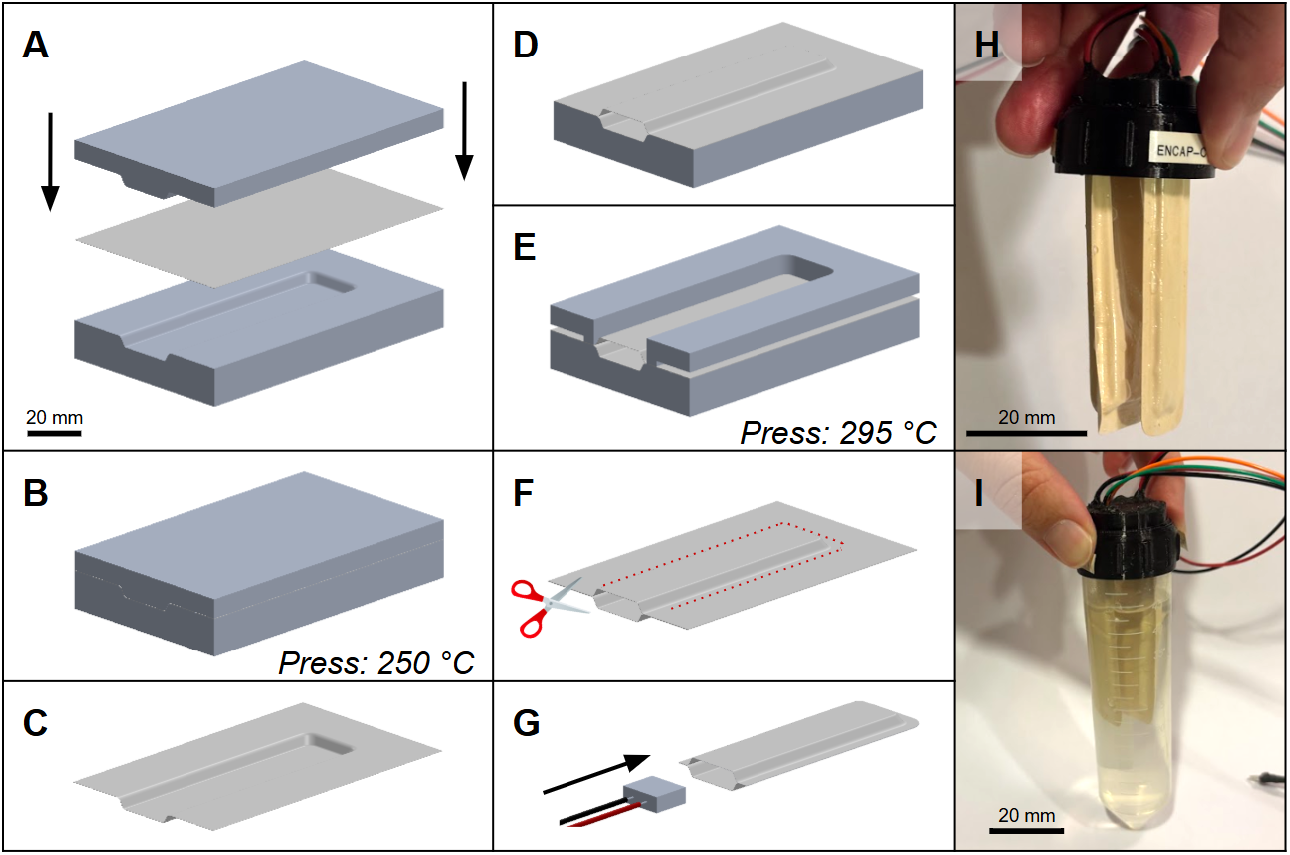
Assembly process for the encapsulated RH sensors, with full description in the text. (A) Alignment of a sheet of LCP into a custom-machined aluminum mold. (B) Closing of the mold after pre-heating, then dwelling at 250 °C for 30-40 minutes, producing (C) one thermoformed LCP sheet. (D) Two thermoformed LCP sheets aligned in the base of the mold, (E) with the sealing fixture placed on top, which is pressed at 295 °C for 30-40 minutes to seal the pocket. (F) The sealed LCP pocket, which is cut to size along the dotted line to produce the pocket shown in (G), which houses an RH sensor. (H) Two sensors mounted into a 3D printed cap with waterproof epoxy. (I) The full assembly screwed onto a 50 mL falcon tube filled with PBS. (A) through (G) are drawn to the same scale.

For each pocket, two pieces of LCP film (LE4277 LFE Liquid Crystal Polymer, Stanford Advanced Materials, Santa Ana, CA, USA) of the desired thickness (25 or 100 μm) were thermoformed to the shape of half of the pocket (3 × 10 × 80 mm and 6 × 20 × 80 mm for capacitive and resistive sensors, respectively, with 30 degree draft angles on walls to prevent excess thinning of the LCP in corners) using a custom machined aluminum mold. Thermoforming was performed at 250 °C for 30-40 minutes in a thermal press (Model 4386, Carver, Wabash, IN, USA), ensuring that the mold and LCP reached temperature prior to fully closing the mold to prevent creasing of the LCP and applying minimal compressive pressure^2^ with the press to prevent thinning of the LCP (Figure 1A-C). After thermoforming, two formed half-pockets were aligned using the base of the mold and a complementary custom machined aluminum sealing fixture was placed on top of the assembly (which applied pressure to the sealing edge of the pocket; Figure 1D-E). Kapton film was used between all LCP and mold/fixture surfaces to prevent sticking during sealing, and hand-molded aluminum foil was inserted into the pocket to ensure it did not collapse during sealing. The assembled fixture was heated to 295 °C in the thermal press with minimal compressive pressure for 30-40 minutes.

After cooling and removing from the fixture, the sealed edges of the pocket were trimmed to a width of approximately 3 mm using precision scissors, and pockets were trimmed to a final length of 80 mm (Figure 1F).

Sensors were inserted into the encapsulation pockets and advanced to the bottom (Figure 1G), then the wires and open end of the pocket were threaded into a 3d printed cap designed to mate with a 50 mL falcon tube. Each cap housed one capacitive and one resistive sensor. The pocket and wire ends were held in place and the pocket opening was sealed off with waterproof epoxy (Ultimate Epoxy, Gorilla, Sharonville, OH, USA; Figure 1H). The cap was screwed onto a 50 mL falcon tube filled with 1x PBS, submerging the encapsulated sensor in the PBS. The final assembly is shown in Figure 1I.

### 3.2 Interdigitated Electrodes

To measure robustness as a flex circuit dielectric, insulated interdigitated electrodes (IDEs) were fabricated using LCP as the insulation material. While submerged in 1x PBS, IDE impedance was monitored.

Interdigitated electrodes were inkjet printed onto LCP film of both 25 and 100 μm thickness using the process described in the following paragraph and illustrated in Figure 2. The IDEs had 0.4 mm finger width and gap, 20 fingers, 5 mm IDE width, and a 90 mm overall device length (including the IDE, traces, and FPC cable). Traces for two IDEs were routed adjacent to each other at the top of the assembly to form a single 4-contact FPC cable.

**Figure 2.**
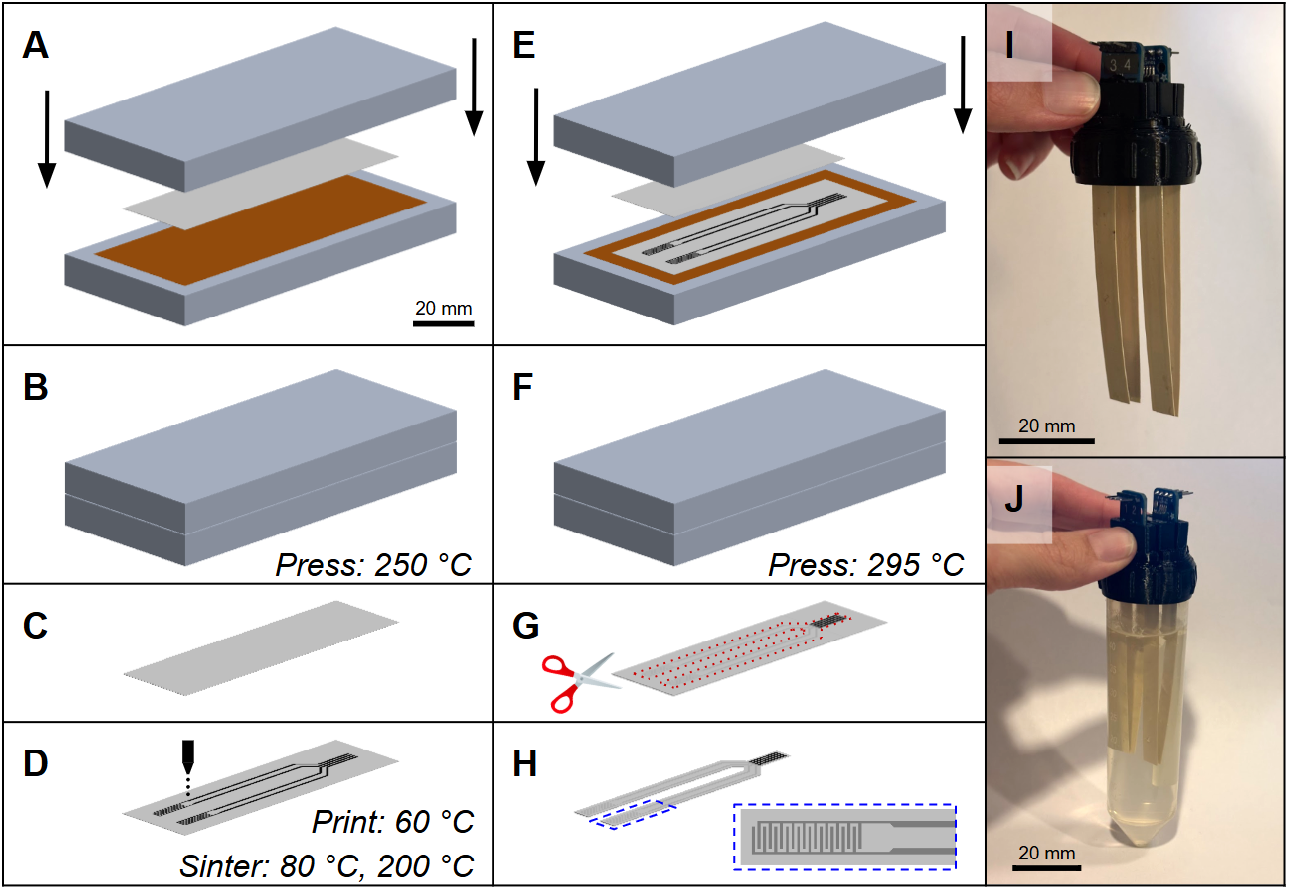
Assembly process for the IDEs, with full description in the text. (A) Placement of a sheet of LCP between two mirror-finish aluminum plates with a Kapton release sheet. (B) Pressing of the LCP sheet at 250 °C for 30-40 minutes, producing (C) one flattened LCP sheet. (D) Two IDEs are printed onto the flattened LCP sheet using silver ink dispensed with an inkjet printer, with the print bed heated to 60 °C. Printed IDEs are sintered at 80 °C, then 200 °C for 15 minutes each. (E) The printed IDEs are placed on a mirror-finish aluminum plate with a second LCP sheet on top of it, leaving the connector end exposed and using Kapton sheets as a release layer. (F) The IDEs are sealed at 295 °C for 30-40 minutes, producing (G) the sealed IDEs, which are cut to size along the dotted line to produce an assembly of two IDEs connected to a single 4-channel FPC cable shown in (H), with the blue dashed inset showing the IDE geometry. (I) Two pairs of IDEs mounted into a 3D printed cap. (J) The full assembly screwed onto a 50 mL falcon tube filled with PBS. (A) through (H) are drawn to the same scale.

For each pair of IDEs, an LCP sheet was flattened between two mirror finish aluminum plates, using Kapton sheets on either side to prevent adhesion to the plates. The assembly was heated to 250 °C in the thermal press with minimal compressive pressure for 30-40 minutes (Figure 2A-C). The flattened sheet was mounted to an inkjet printer (DMP-2831, Fujifilm Dimatix Inc, Santa Clara, CA, USA) print bed using vacuum and kapton tape, and the IDE was printed with silver ink (DM-SIJ-3206, Dycotek, Wiltshire, UK) with 20 layers using a 2.4 pL/drop cartridge (Samba, Fujifilm Dimatix Inc, Santa Clara, CA, USA) and allowing 5-10 minutes of drying time at 60 °C between layers. The ink was dried and sintered at 80 °C and 200 °C for 15 minutes each (Figure 2D). The printed IDE was placed on top of a mirror finish aluminum plate with a Kapton sheet to prevent adhesion to the plate, and a second LCP sheet of the same thickness was placed on top of the IDE, leaving the FPC connector area exposed. A second Kapton sheet and mirror finish plate was placed on top, and the assembled fixture was heated to 295 °C in the thermal press with minimal compressive pressure for 30-40 minutes (Figure 2E-F). After cooling and removing from the fixture, the laminated IDEs were trimmed to approximately 1 mm from the printed edge using precision scissors (Figure 2G-H).

Each pair of IDEs were connected to a 4-contact FPC connector breakout board (3575, Adafruit, Brooklyn, NY, USA), using 1-2 pieces of 100 μm thick, 5 × 10 mm LCP inserted behind the IDE cable to thicken it sufficiently to engage the connector pins. Connected IDEs were threaded into a 3d printed cap designed to mate with a 50 mL falcon tube (Figure 2I). Each cap housed two pairs of IDEs. The opening in the cap was sealed and the breakout boards were held in place with waterproof epoxy. The cap was screwed onto a 50 mL falcon tube filled with 1x PBS, submerging the IDEs in the PBS. The final assembly is shown in Figure 2J.

### 3.3 Accelerated Lifetime Test Setup

Samples mounted in falcon tubes were placed in a dry bath (2510-1104, USA Scientific, Ocala, FL, USA) targeting 67 °C, equating to 8x acceleration at body temperature per the arrhenius equation. Actual temperature of the dry bath was continuously measured and used for more accurate accelerated time calculations for all samples. For sample groups where multiple temperature sensors were used, the average temperature across all sensors at a given time point was used. PBS was replaced or replenished as needed due to evaporation (approximately once per month for the first six months, then reduced to approximately once per four months after adding mineral oil on top of the PBS to prevent evaporation).

The capacitive humidity sensors had an I^2^C output which was monitored with an arduino (UNO Rev 3, Arduino, Monza, Italy). The sensor I^2^C address was not changeable, so custom hardware and an I^2^C scanner was used to read from all four sensors with a single arduino I/O. RH and temperature were read directly from the sensor; temperature was averaged across all working sensors.

Resistive humidity sensors were measured with an LCR meter (LCX100, Rohde & Schwarz, Columbia, MD, USA) at 1 kHz and 1 V excitation. The sensor was rated to operation temperatures of 70 °C, however the datasheet provided calibration from impedance to RH at discrete operating temperatures up to only 45 °C (which required extrapolation at the desired operating temperature of approximately 67 °C). Temperature readings were taken from the average reading of the capacitive sensors.

IDE impedance was measured with the same LCR meter from 10 Hz to 10 kHz at 25 mV excitation. Impedance at 1 kHz was used for longevity comparison in this work, but all data are publicly available on GitHub. Temperature was measured with a thermocouple in the dry bath, and temperature offset was calibrated by measuring PBS temperature with a thermocouple one week into the experiment.

For the first 30-45 days, measurements were taken manually at 1-3 day intervals, after which an automated setup was implemented. A block diagram of the full automated accelerated lifetime test setup is shown in Figure 3. Resistive humidity sensors and IDEs were connected to the LCR meter through a multiplexer (7002 Switch System equipped with 7011-C multiplexer cards, Keithley, Solon, OH, USA) so that a single sample could be connected and measured at a time programmatically. Control of the multiplexer, LCR meter, and arduino were all integrated into a single python script which would collect, log, and plot data every 12 hours. Python and arduino scripts are publicly available on GitHub.

**Figure 3.**
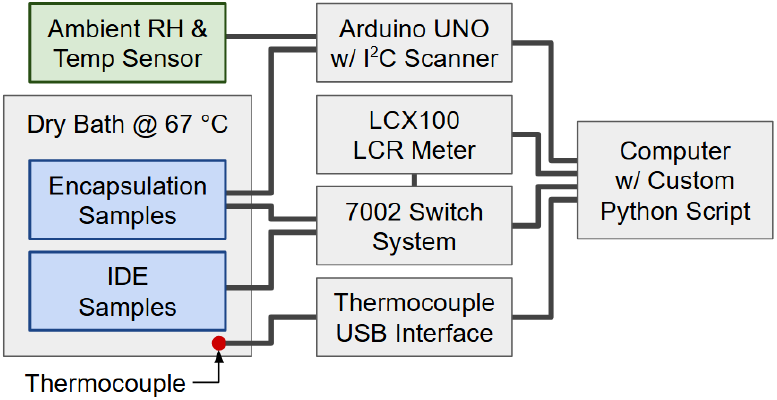
Block diagram of the accelerated lifetime test setup

## 4. Results

### 4.1 Experimental Notes

Two early experimental issues impacted portions of the data, however both were relatively minor and did not impact the ability to draw conclusions from the full dataset. These problems are described briefly here, and the full dataset is included below, with any relevant effects noted as appropriate.

First, during sample preparation, only one type of LCP was available. Typical LCP fabrication requires two formulations of LCP with different melting temperatures, with the lower temperature LCP acting as an adhesive layer or coverlay. Due to this limitation, several early samples experienced engineering failures (thinning or pinholes due to over-softening of LCP during lamination). Despite these issues, these data are presented here because the low temperature LCP was unavailable for several months and the collected data from the good samples demonstrates excellent performance of LCP under chronic soaking conditions.

Second, data collected during the first half of the experiment (33.5 weeks in the encapsulation group, 32 weeks in the IDE group) have significantly higher noise than data collected in the latter half of the experiment. This was due to an external noise source located near the experimental setup that was removed. This effect is more pronounced in the IDEs because these samples were connected to the measurement equipment via longer, unshielded wires as compared to the RH sensors.

### 4.2 Encapsulation Samples

The two resistive RH sensors encapsulated in 25 μm LCP experienced pinhole failures, flooding the pocket with PBS within several hours of starting the experiment. This was likely due to over thinning of the LCP when producing the thicker pocket for the resistive sensor in combination with re-heating the pocket to its melting temperature when sealing the two LCP pieces together. The capacitive sensors had thinner pockets, decreasing risk of over-thinning of the LCP, and the 100 μm thick LCP had sufficient thickness to survive thermoforming at either size. Also, a software error incorrectly indicated that one capacitive RH sensor had failed, and so it was removed from the study; this device was later repurposed as the ambient sensor. After failures and troubleshooting, one capacitive sensor in the 25 μm group and all four sensors in the 100 μm group remained under test.

RH versus accelerated time for all surviving encapsulated samples is shown in Figure 4. The RH inside the encapsulation samples, as measured with the capacitive sensors, remained very low and stable over 61 weeks at an average temperature of 64.9 °C. This equates to an equivalent lifetime of 8.1 years at 37 °C. Minor fluctuations in RH generally mirror changes in sample temperature and ambient RH and temperature. This is to be expected: RH is defined as the ratio of vapor pressure to saturation vapor pressure, and since the latter increases with higher temperature, the RH of a sample will decrease with higher temperature. These fluctuations are particularly apparent with changes, such as the local maximum sample temperature at 11 weeks corresponding to a local minimum in RH, or the global maximum ambient humidity at 59 weeks corresponding to a small visible peak in RH. For all capacitive sensors, baseline RH was in the range of 8-15% and all measurements were in the range of 5-16% (excluding the initial two measurements during temperature stabilization).

**Figure 4.**
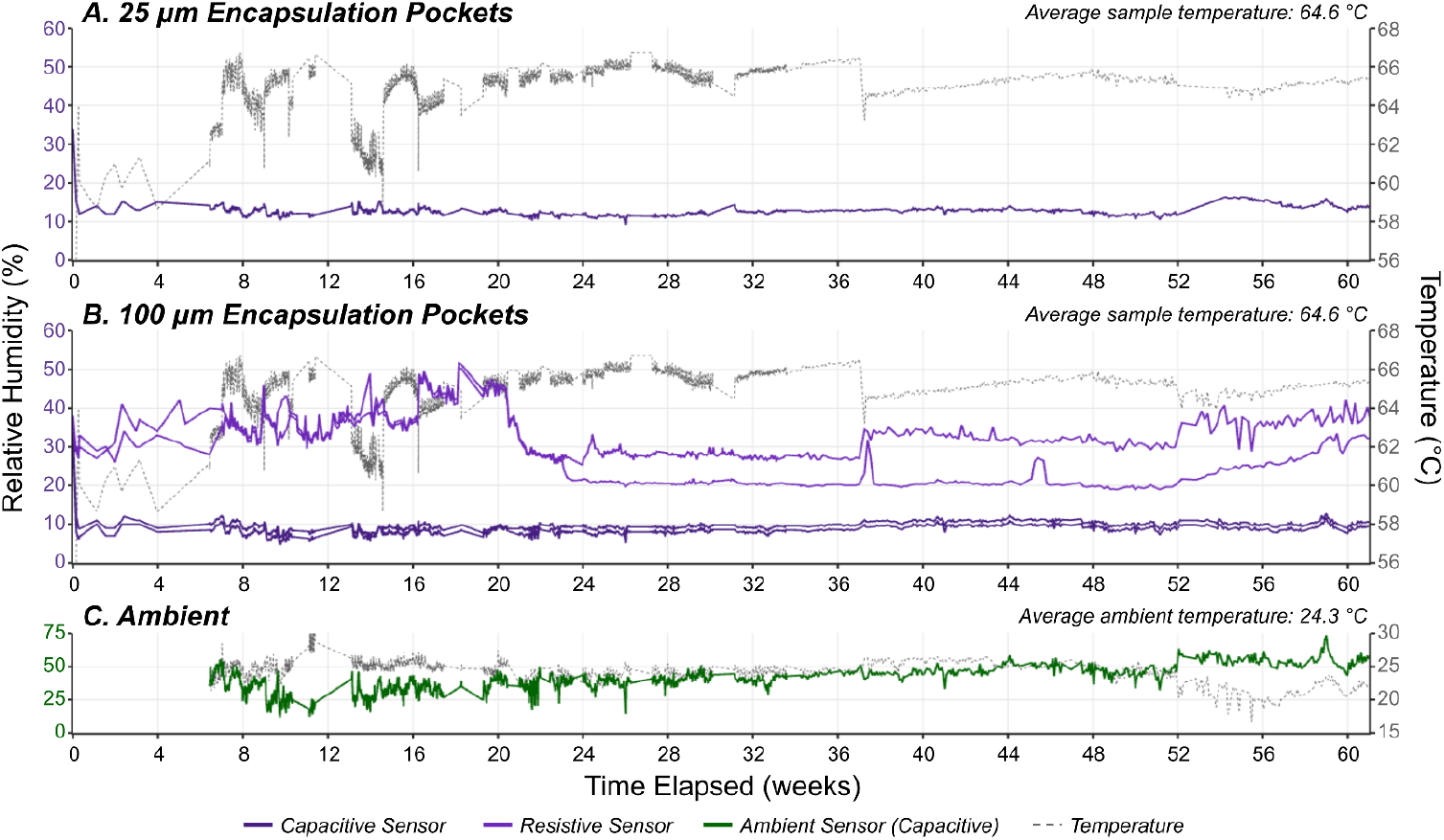
Relative humidity and temperature versus accelerated time for relative humidity sensors encapsulated in (A) 25 μm and (B) 100 μm thick LCP. The dark and light purple lines represent relative humidity of the capacitive and resistive sensors, respectively. (C) shows the ambient relative humidity in green. Gray dotted lines in all plots show temperature of the given sample (the PBS bath or ambient). The change in signal variance at 33.5 weeks is attributed to the removal of an external noise source, as described in section 4.1.

RH measured with the resistive sensors (inside only 100 μm thick LCP, owing to pinholes created during thermoforming of the thick pockets with the 25 μm LCP) had a higher value and more fluctuations, with baseline values of 29-30% and a full range of 19-51% (excluding the initial two measurements during temperature stabilization). The higher variance and offset in this data stems from two likely sources. First is the higher sensitivity of these sensors to temperature fluctuation. Similarly to the capacitive sensors, fluctuations in the resistive RH data mirror fluctuations in sample temperature and ambient temperature and RH, but with an apparent greater impact of temperature. This is primarily visible during temperature changes, such as the sample temperature decrease seen at 37 weeks corresponding to an increase in RH read by one resistive sensor, or the ambient temperature decrease between 52 and 56 weeks corresponding to an increase in RH read by both resistive sensors. Second is the error introduced during conversion of the resistance readings to RH. The manufacturer-provided datasheet included calibration curves at discrete temperature profiles up to 45 °C, so a conversion relating to variable resistance and temperature to RH was extrapolated from these data. As such, these sensors are primarily included not for their absolute values, but as secondary means of detecting RH changes and, in particular, catastrophic failures, such as delamination of the LCP leading to flooding of the encapsulation pocket should they occur in the future.

### 4.3 Interdigitated Electrodes

Impedance magnitude change from baseline versus accelerated time for all IDEs is shown in Figure 5. Raw impedance values are not reported here because they varied significantly between IDEs due to the variability in the inkjet printing process, however raw impedance data is available on GitHub.

**Figure 5.**
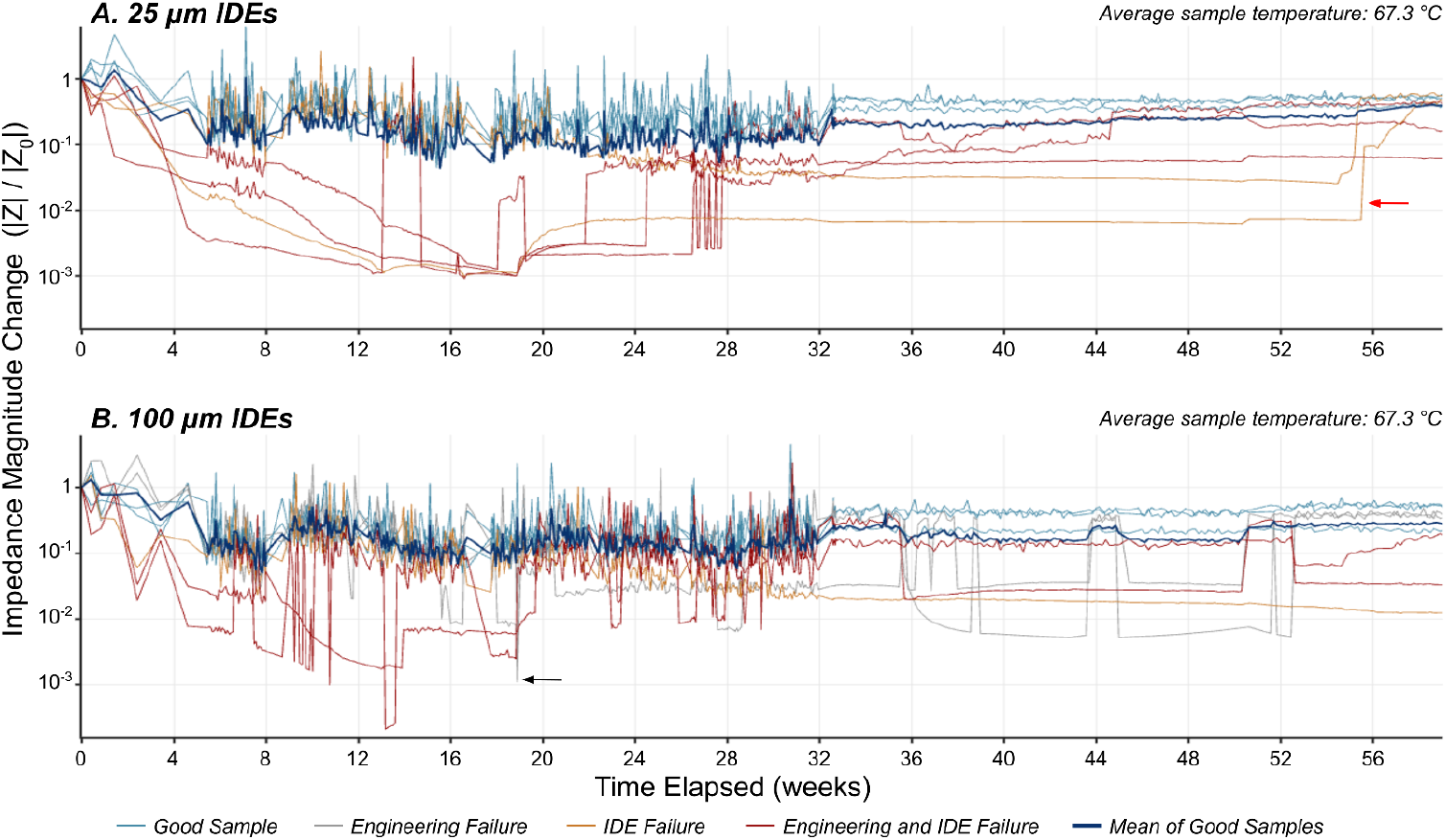
Impedance magnitude change from baseline (|Z| / |Z|0) versus accelerated time for IDEs laminated in (A) 25 μm and (B) 100 μm thick LCP. The light blue lines represent good samples, the gray lines represent samples which had engineering failures, the orange lines represent samples which had IDE failures, and the red lines represent samples which had both engineering and IDE failures (as defined in the text of section 4.3) The dark blue solid lines represent the mean value of the good samples. The black arrow points to one example of a transient, high magnitude signal change indicative of a connection issue, classified as an “:engineering failure”:. The red arrow points to one example of a high magnitude signal change likely indicative of metal degradation on the sensor. The change in signal variance at 32 weeks is attributed to the removal of an external noise source, as described in section 4.1.

Several IDE samples in both the 25 μm and 100 μm LCP groups exhibited marked, rapid drops and increases in impedance, such as the sample which went from |Z| / |Z|_0_ = 0.7 to 0.001 to 0.13 in three subsequent measurements taken 12 hours apart, marked by a black arrow in Figure 5B. These are assumed to be due to transient connector issues. Rapid decreases in impedance could be indicators of connector issues (shorting of connector contacts due to moisture), catastrophic IDE failure (exposing the IDE to PBS), or some other electrode issue unrelated to the LCP insulation, while rapid increases in impedance could be an indicator of disconnection from the connector. However, catastrophic IDE or connector failures would not exhibit recovery back to or near the baseline value, so samples with two or more rapid impedance increases within subsequent samples were categorized as engineering failures, rather than failures of the LCP itself. These data are shown in gray in Figure 5.

In contrast, IDE failures were characterized by gradual impedance decrease from the baseline value over the course of several weeks or months. Specifically, samples whose impedance magnitude change from baseline saw a gradual decrease to a value below 0.001 within the first 12 weeks, or to a value less than 0.01 at the time of writing (59 weeks) were characterized as IDE failures. These are shown as orange lines Figure 5. Samples which could be classified as both engineering and IDE failures are shown in red in Figure 5. Some of these samples saw subsequent recovery back towards their baseline impedance, such as the sample indicated by the red arrow in Figure 5A. Although the cause of this is not confirmed, this could be due to causes such as connector issues or degradation of the printed metal after chronic exposure to moisture.

The remainder of the IDEs (i.e. those which did not experience either of the failures described above) are classified as functional, and their data is shown in blue in Figure 5. The impedance magnitude of functional IDEs decreased slightly over the first several weeks, then has remained stable for the remainder of the 59 weeks of testing to date, with an average temperature of 67.5 °C over the entire testing period. This equates to an equivalent lifetime of 9.4 years at 37 °C. Impedance changes from baseline measured in the past 6 weeks (equivalent to 1 year at 37 °C) range from 0.21-0.70 (mean values 0.48 and 0.40 for 25 μm and 100 μm sample groups, respectively). This likely reflects some saturation of the LCP with water, as expected, since absorbed water would slightly decrease the impedance between IDE contacts, but does not indicate chronic failure of the material. Similarly, no delamination of LCP was visible on any samples. Finally, there was no noticeable difference between the 25 μm and 100 μm sample groups, though sample sizes were not large enough to perform statistical analysis.

## 5. Discussion

These results confirm good moisture barrier performance of LCP and demonstrate those properties *in vitro* over 8+ equivalent years of implanted time. These results are consistent with the known moisture barrier properties of LCP, such as WVTR and moisture absorption, yet no prior studies have evaluated the long-term LCP performance without confounding conditions, such as stimulating electrode performance. Note that the data presented here represent a snapshot of progress and the study remains ongoing, with all data publicly available on GitHub.

### 5.1 Encapsulation Samples

The data reported here show stable performance of LCP as a moisture barrier material for implanted electronics, with an estimated equivalent lifetime of more than 8.1 years at 37 °C. Fick’s first law of diffusion says that water will diffuse into a sample with rate proportional to the concentration gradient and a material-specific diffusion coefficient and inversely proportional to the distance (LCP thickness). Thus, the RH measurements shown here indicate that LCP’s WVTR is very low, as expected, yielding undetectable changes in RH. Continued testing will eventually show a slow rise in RH which will reveal the lifetime of LCP as an encapsulant material.

Minor fluctuations in RH data corresponding to changes in ambient humidity and temperature stem from two likely causes: (1) that the air temperature around the experimental setup impacted the temperature of the samples and resulting RH inside the tube and inside the pocket, and (2) that there was a leak in the seal at the top of the encapsulation pocket, allowing air exchange between the ambient air and the inside of the pocket. While care was taken to seal the top of the encapsulation pockets and prevent the exchange of air, a perfect seal was likely not made, and both explanations are possible. Resistive sensor measurements are more sensitive to temperature data, supporting the first cause, while some capacitive sensor fluctuations correspond directly to ambient RH fluctuations (such as the peak at 59 weeks), supporting the second cause. The current experimental setup does not allow for a conclusive test to quantitatively determine the impacts of each; a future experiment to avoid these effects would include a wireless humidity sensor that could be enclosed in LCP on all sides and fully submerged in PBS during testing.

### 5.2 Interdigitated Electrodes

Although the impedance values of the IDEs classified in the functional group have decreased from their initial values, they have remained stable from weeks 32 to 59. This suggests that the LCP absorbed and saturated with fluid during some initial period of time, then reached an equilibrium where it can hold no additional fluid. This is expected per Fick’s first law. Due to the external noise source present during the first 32 weeks of the experiment, however, time to reach saturation cannot be accurately measured, although it appears to fall somewhere in the range of 5-16 weeks for both sample groups. These IDE resistance changes can be used to calculate the expected crosstalk in implanted circuits due to moisture ingress, useful for designing circuits to accommodate for moisture-related shunt resistance.

Despite these encouraging results, engineering failures were also observed. The early failures in encapsulation are presumed to be due to pinhole failures, which lead to flooding of the encapsulation pockets. These failures should be preventable with straightforward engineering fixes such as the use of thicker LCP when larger pockets are required

Engineering failures such as delamination, equipment failure, connector failures should appear as rapid changes in impedance: delamination would cause a shorting of the two IDE contacts and a rapid decrease in impedance, and equipment or connector failures would be seen as intermittently, rapid decreases and increases in impedance. The latter was seen in 4/16 samples. Slower IDE failures were also seen in 6/16 of the devices, as evidenced by gradual impedance decrease. The cause of subsequent recovery in impedance in some samples characterized as IDE failures is unknown, but hypothesized to be due to connector issues or degradation of the printed metal IDEs.

Although the root causes of these failures are not confirmed, visual inspection confirms no bulk delamination. In contrast, the most common microfabrication and encapsulation polymers (polyimide and Parylene C) commonly suffer from delamination failures during chronic use [12,13]. The fact that there is no delamination observed in any samples underscores a key advantage compared to these more widely used materials.

## 6. Conclusion

LCP has been demonstrated as a good material for both encapsulation of electronics and construction of flex circuits for use in implanted medical devices or other applications requiring long term moisture barriers. Here, LCP is demonstrated to maintain moisture resistance over more than 8.1 years of simulated implantation (59+ weeks *in vitro*, thermally accelerated at 7-8x), with experiments ongoing. While some samples saw early failures, these can be attributed to causes that can be solved with further process development. Long term success of the remaining, working samples indicate that these failures can be attributed to yield loss during fabrication and are not a function of material failure.

In working samples, there was no noticeable difference between the 25 μm and 100 μm sample groups, however sample sizes were not large enough for statistical analysis. In the encapsulation sample group, however, yield loss was significantly higher in the 25 μm group due to higher risk of material thinning during the thermoforming process. This indicates that, when fabricated correctly, LCP serves as a reliable moisture barrier down to a thickness of 25 μm.

## 7. Author Contributions

B.T. designed the experiments, fabricated the samples, conducted the testing, analyzed the data, and developed automation for the experimental workflow. C.P. and E.A. contributed to early equipment setup and development of the automation framework. E.A. additionally provided guidance on experimental design and supported early-stage work. P.S. reviewed the data and edited the manuscript. P.S. and M.C. provided overall support for the project.

## 8. Supplementary Data

All data presented here plus additional raw data, Arduino code for capacitive sensor readings, and Python automation scripts are available on GitHub at the link below. At the time of writing, experiments are ongoing and the raw data and plots are updated after every measurement (with measurement intervals decreased to 24 hours).

GitHub Repository: https://github.com/brianna-thielen/Lifetime-Testing

## Footnotes

1 WVTR value calculated at a film thickness of 10 μm.

2 In all instances, “:minimal compressive pressure”: indicates that pressure was not measured as it was below the gauge resolution of the thermal press (600 lb over 860 and 900 mm^2^ area for capacitive and resistive pockets, respectively), however it was just past where the press engaged and likely in the 10s of lbs of force.

## References

1. Bigham KJ. LCP Introduction to Liquid Crystal Polymers. Zeus Technical Papers. 2018.

2. Lee SW, Min KS, Jeong J, Kim J, Kim SJ. Monolithic Encapsulation of Implantable Neuroprosthetic Devices Using Liquid Crystal Polymers. IEEE Trans Biomed Eng. 2011; 58(8): 2255–2263. 10.1109/TBME.2011.2136341.

3. Pak A, Nanbakhsh K, Hölck O, Ritasalo R, Sousa M, Van Gompel M, Pahl B, Wilson J, Kallmayer C, Giagka V. Thin Film Encapsulation for LCP-Based Flexible Bioelectronic Implants: Comparison of Different Coating Materials Using Test Methodologies for Life-Time Estimation. Micromachines. 2022; 13(4): 544. 10.3390/mi13040544.

4. Overview of materials for Liquid Crystal Polymer (LCP), Unfilled. MatWeb. Accessed 14 January 2026.

5. LE4277 LFE Liquid Crystal Polymer Test Report. Stanford Advanced Materials. Delivered 28 October 2024.

6. Kaiser A, Bee CM, Dupuis F, von Metzen R, Rueß K, Matej P, Herbort C, Holl B, Löffler S, Bauböck G, Fritz K-H. Thin Film Based LCP Multi-Layer Circuits: Manufacturing Technology and Characterization. IEEE EMPC 2015; 1–6.

7. Ahn S-H, Koh CS, Park M, Jun SB, Chang JW, Kim SJ, Jung AHH, Jeong J. Liquid Crystal Polymer-Based Miniaturized Fully Implantable Deep Brain Stimulator. Polymers. 2023; 15(22): 4439. 10.3390/polym15224439.

8. Rachinskiy I, Wong L, Chiang C-H, Wang C, Trumpis M, Ogren JI, Hu Z, McLaughlin B, Viventi J. High-Density, Actively Multiplexed μECoG Array on Reinforced Silicone Substrate. Front Nanotechnol. 2022; 4: 837328. 10.3389/fnano.2022.837328.

9. Sun T, Park W-T, Cheng M-Y, An J-Z, Xue R-F, Tan K-L. Implantable Polyimide Cable for Multichannel High-Data-Rate Neural Recording Microsystems. IEEE Trans Biomed Eng. 2011; 59(2): 390–399. 10.1109/TBME.2011.2173343.

10. Buchwalder S, Nicolier C, Hersberger M, Bourgeois F, Hogg A, Burger J. Development of a Water Transmission Rate (WTR) Measurement System for Implantable Barrier Coatings. Polymers. 2023; 15(11): 2557. 10.3390/polym15112557.

11. Woods V, Trumpis M, Bent B, Palopoli-Trojani K, Chiang C-H, Wang C, Yu C, Insanally MN, Froemke RC, Viventi J. Long-term recording reliability of liquid crystal polymer μECoG arrays. J Neural Eng. 2018; 15(6): 066024. 10.1088/1741-2552/aae39d.

12. Ortigoza-Diaz J, Scholten K, Meng E. Characterization and Modification of Adhesion in Dry and Wet Environments in Thin-Film Parylene Systems. J Microelectromech Syst. 2018; 27(5): 874–885. 10.1109/JMEMS.2018.2854636.

13. Schander A, Gancz J M, Tintelott M, Lang W. Towards Long-Term Stable Polyimide-Based Flexible Electrical Insulation for Chronically Implanted Neural Electrodes. Micromachines. 2021; 12(11): 1279. 10.3390/mi12111279.

